# Orchestrating and sharing large multimodal data for transparent and reproducible research

**DOI:** 10.1101/2020.09.18.303842

**Authors:** Anthony Mammoliti, Petr Smirnov, Minoru Nakano, Zhaleh Safikhani, Christopher Eeles, Heewon Seo, Sisira Kadambat Nair, Arvind S. Mer, Chantal Ho, Gangesh Beri, Rebecca Kusko, MAQC Society, Benjamin Haibe-Kains

## Abstract

Reproducibility is essential to open science, as there is limited relevance for findings that can not be reproduced by independent research groups, regardless of its validity. It is therefore crucial for scientists to describe their experiments in sufficient detail so they can be reproduced, scrutinized, challenged, and built upon. However, the intrinsic complexity and continuous growth of biomedical data makes it increasingly difficult to process, analyze, and share with the community in a FAIR (findable, accessible, interoperable, and reusable) manner. To overcome these issues, we created a cloud-based platform called ORCESTRA (<u>orcestra.ca</u>), which provides a flexible framework for the reproducible processing of multimodal biomedical data. It enables processing of clinical, genomic and perturbation profiles of cancer samples through automated processing pipelines that are user-customizable. ORCESTRA creates integrated and fully documented data objects with persistent identifiers (DOI) and manages multiple dataset versions, which can be shared for future studies.

## INTRODUCTION

The demand for large volumes of multimodal biomedical data has grown drastically, partially due to active research in personalized medicine, and further understanding diseases^1–3^. This shift has made reproducing research findings much more challenging because of the need to ensure the use of adequate data handling methods, resulting in the validity and relevance of studies to be questioned^4,5^. Even though sharing of data immensely helps in reproducing study results^6^, current sharing practices are inadequate with respect to the size of data and corresponding infrastructure requirements for transfer and storage^2,7^. As computational processing required to process biomedical data is becoming increasingly complex^3^, expertise is now needed for building the tools and workflows for this large-scale handling^1,2^. There have been multiple community efforts in creating standardized workflow languages, such as the Common Workflow Language (CWL) and the Workflow Definition Language (WDL), along with associated workflow management systems such as Snakemake^8^ and Nextflow^9^, in order to promote reproducibility^10,11^. However, a steep learning curve is encountered for these programming-heavy solutions, in comparison to user-friendly data processing platforms like Galaxy, which provide both storage and compute resources, but have limited features and scalability^12–14^. Therefore, there is a dire need for reproducible and transparent solutions for processing and analyzing large multimodal data that are scalable, while providing full data provenance.

Biomedical data can expand into a plethora of data types such as *in vitro* and *in vivo* pharmacogenomics, toxicogenomics, radiogenomics and clinical genomics. These data are a prime example of multimodal biomedical data with a long history of sharing in the field of biomarker discovery. Preclinical pharmacogenomics involves the use of a genome-wide association approach to identify correlations between compound/treatment response and molecular profiling, such as gene expression^15–17^. In addition, omics technologies have also been utilized in toxicological profiling for identifying the effect of compound toxicity on humans^18^, and in radiogenomics data to uncover genomic correlates of radiation response^19^. These rich preclinical data are often combined with clinical genomics data generated over the past decades^20^ with the aim to test whether preclinical biomarkers can be translated in clinical settings to ultimately improve patient care. Given the diversity of human diseases and therapies, researchers can hardly rely on a single dataset and benefit from collecting as much data as possible from all possible sources, calling for better sharing of data that are highly standardized and processed in a transparent and reproducible way.

The generation of large volumes of data has led to a sharing paradigm in the research community, where data is more accessible and open for public use. For studies to be reproduced and investigated for integrity and generalization by other researchers, the sharing of raw and processed data is crucial. However, providing access to data is not enough to achieve full reproducibility, as the shared data must be findable, accessible, interoperable and reusable, as outlined in the FAIR data principles^21^. These foundational principles include providing rich metadata that is detail-oriented, including a persistent unique-identifier (Findability), accessing (meta)data with authentication and the unique-identifier using a communications protocol (Accessibility), assigning (meta)data with a commonly understood format/language (Interoperability), and achieving data provenance with an accessible usage-license (Reusability). The Massive Analysis and Quality Control (MAQC) Society^22^ has been established to promote the use of a community-agreed standard for sharing multimodal biomedical data in order to achieve reproducibility in the field, such as through the FAIR principles. Therefore, when translated into practice, these principles would promote the reproducible and transparent handling and sharing of data and code, which would allow researchers to utilize and build from each other’s work and accelerate new discoveries.

In order to address these issues, we developed ORCESTRA (orcestra.ca) a cloud-based platform that provides a transparent, reproducible, and flexible computational framework for processing and sharing large multimodal biomedical data. The ORCESTRA platform orchestrates data processing pipelines in order to curate customized, versioned and fully documented data objects, which can be extended to a multitude of data types. This includes 11 pharmacogenomics (in vitro), 3 toxicogenomics, 1 xenographic pharmacogenomics (in vivo), 1 clinical genomics, and 1 radiogenomics data objects that can be explored for a wide range of analyses. ORCESTRA is publicly accessible via orcestra.ca.

## RESULTS

The increasing utilization and demand for big data have resulted in the need for effective data orchestration^23^, which is a process that involves organizing, gathering, and coordinating the distribution of data from multiple locations across compute resources with specific processing requirements. An ideal orchestration platform for handling large-scale heterogeneous data would consist of the following: (1) a defined workflow; (2) a programming model/framework^23^, and (3) broad availability of a compute infrastructure. At the workflow level, data from different sources/lineages, including data that are not static, must be effectively managed through the definition of workflow components (tasks) that interact and rely on one another^23^. Moreover, a programming model should be utilized for the workflow components responsible for handling the respective data (static and dynamic), such as a batch processing model (e.g., MapReduce)^23^. Lastly, the utilization of a scalable compute environment, such as academic and commercial cloud computing platforms, would allow for the management and processing of big data, providing the necessary compute and storage resources, ability to transfer data, and monitoring of executed workflows and respective components/tasks, further enabling tracking data provenance. There exist multiple orchestration tools that are currently being used for the storage, processing, and sharing of genomic data, namely Pachyderm, DNAnexus, Databricks, and Lifebit (Table 1). We opted for Pachyderm, an open-source orchestration tool for multi-stage language-agnostic data processing pipelines, maintaining complete reproducibility and provenance through the use of Kubernetes, as it provides the following functionalities:

**Table 1.**
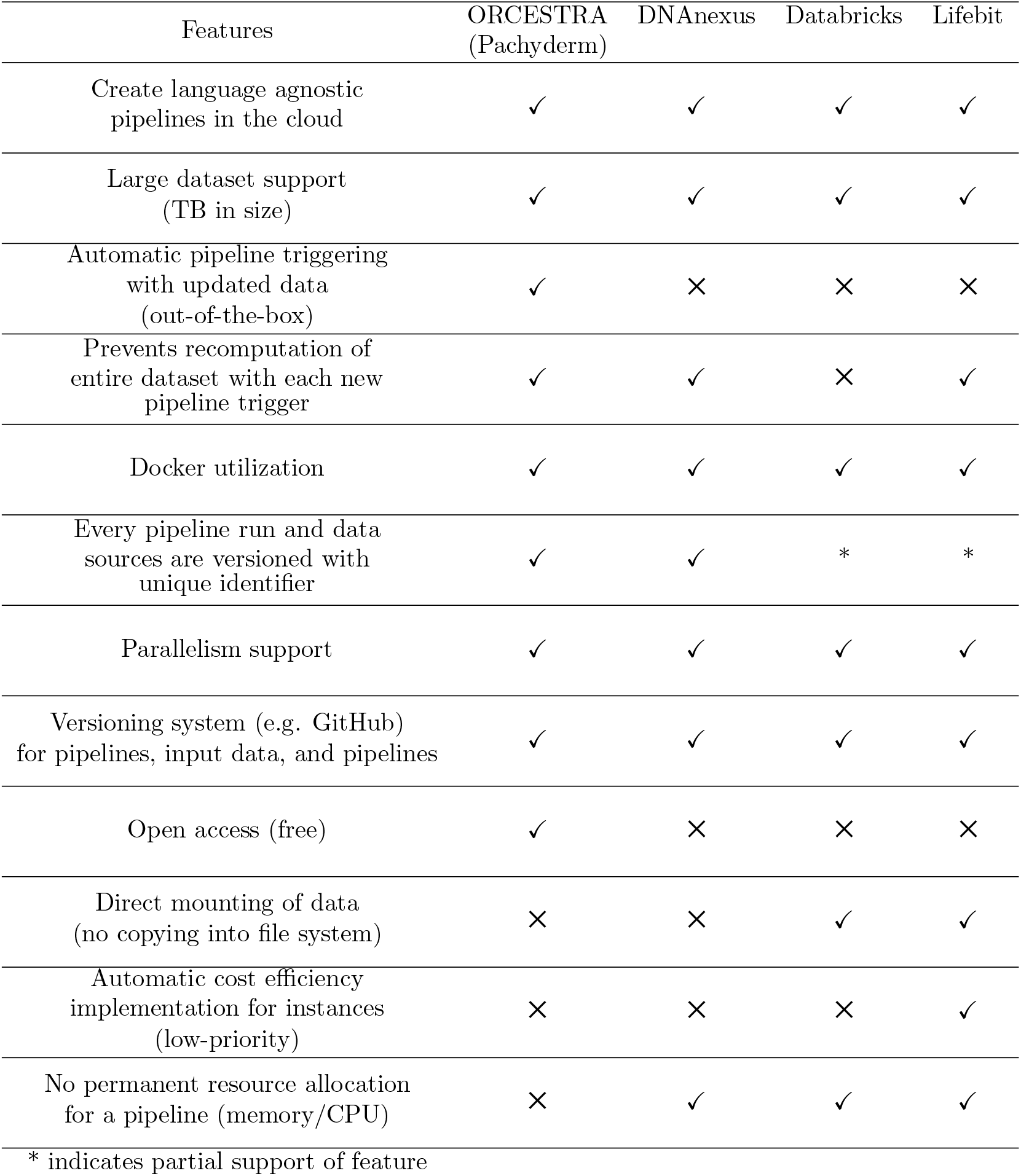
Data processing platforms and their respective features for handling multimodal data.

| Features | ORCESTRA<br>(Pachyderm) | DNAnexus | Databricks | Lifebit |
| --- | --- | --- | --- | --- |
| Create language agnostic pipelines in the cloud | ✓ | ✓ | ✓ | ✓ |
| Large dataset support (TB in size) | ✓ | ✓ | ✓ | ✓ |
| Automatic pipeline triggering with updated data (out-of-the-box) | ✓ | × | × | × |
| Prevents recomputation of entire dataset with each new pipeline trigger | ✓ | ✓ | × | ✓ |
| Docker utilization | ✓ | ✓ | ✓ | ✓ |
| Every pipeline run and data sources are versioned with unique identifier | ✓ | ✓ | * | * |
| Parallelism support | ✓ | ✓ | ✓ | ✓ |
| Versioning system (e.g. GitHub) for pipelines, input data, and pipelines | ✓ | ✓ | ✓ | ✓ |
| Open access (free) | ✓ | × | × | × |
| Direct mounting of data (no copying into file system) | × | × | ✓ | ✓ |
| Automatic cost efficiency implementation for instances (low-priority) | × | × | × | ✓ |
| No permanent resource allocation for a pipeline (memory/CPU) | × | ✓ | ✓ | ✓ |
| * indicates partial support of feature |  |  |  |  |

### Programming language

Pachyderm supports creating and deploying language-agnostic pipelines across on-premise or cloud infrastructures, a feature also supported by DNAnexus, Databricks, and Lifebit.

### Large dataset support

Users can upload and process large datasets through the use of the Pachyderm file system (PFS), where the data is exposed in its respective container for utilization in pipelines, while being placed in an object storage (e.g., Azure Blob, AWS bucket).

### Automatic pipeline triggering

Reproducibility and provenance are guaranteed via automatic pipeline triggering when existing data are modified or newly added, which results in the generation of new versions of an output data object. However, because automatic triggering requires the state of each pod within the Kubernetes cluster to be saved, there is a permanent allocation of CPU/RAM for each pod (and therefore each pipeline), which requires a user to create a cluster with potentially costly resources. The other platforms do not require permanent allocation of resources, as for example Lifebit allows users to spin up instances on demand to meet the computational requirements for a given pipeline.

### Reprocessing

A feature that is found in Pachyderm, DNANexus, and Lifebit is the prevention of recomputation for each pipeline trigger, which comes in handy when a pipeline contains processed raw data that does not need to be reprocessed if there is a change in metadata such as an annotation file.

### Docker utilization

Each pipeline can be equipped with a Docker image connected to Docker Hub for running various toolkits, which allows for simplistic pipeline updating when there are future updates to any component of the Docker image. Docker usage is also translated across the other platforms as well.

### Versioning of data and pipelines with unique identifiers

Each commit, an operation for submitting and tracking changes to a data source, is supplied with a unique identifier, which is updated with each new commit (parent-child system). This allows users to track different versions of a pipeline and dataset with ease. However, with Databricks and Lifebit, this feature is partially supported, as not every pipeline and respective input/output file(s) are provided with a unique identifier, even when data is updated through commits.

### Parallelism support

A pipeline can be parallelized via a constant or coefficient strategy in Pachyderm using workers, which is useful for workloads with large computational requirements. When a constant is set Pachyderm will create the exact number of workers specified (e.g., constant: 5; 5 workers), that will parallelize across nodes in the cluster. Coefficient will result in Pachyderm creating a number of workers based on the number of nodes available (multiple of nodes), which will also specify the number of workers per node (e.g., coefficient: 2.0; 20 nodes; 40 workers; 2.0 workers per node). The other platforms also support parallelization, including automatic parallelization of samples across instances.

### Data versioning system

Pachyderm provides direct GitHub integration for data versioning, which enables users to track changes at the file-level and submit updates to Pachyderm through commits triggered through webhooks on GitHub. In addition, this also provides users with the ability to publicly view, track, and share all updates made to a pipeline or file connected to Pachyderm with ease.

### Open access

Pachyderm provides a free and open-source version of the tool that contains all the functionalities required to develop a platform ensuring transparent and reproducible processing of multimodal data.

Despite these advantages, the choice of Pachyderm is not without compromises. We list below the functionalities that Pachyderm is lacking but would have been beneficial to develop our platform:

### Direct mounting of data

The current version of Pachyderm does not allow for direct mounting of data from a cloud storage system (e.g., bucket) to a Pachyderm repository. Data must be transferred to the tools own file system, resulting in essentially an additional copy of the data within a cloud environment. Databricks and Lifebit enable decreasing computation time and cost by not copying data into a file system for it to be used by the platform. This is important when large data sizes will be used in an analysis, which allows a user to simply store their data in a bucket/blob storage account, and mount it to the platform of interest, giving the user the ability to also use the data with other platforms or cloud services without having to repeatedly copy it in an inefficient manner.

### Cost efficiency

Pachyderm utilizes VM’s through a Kubernetes cluster if deployment on a cloud environment, which are costly to keep running indefinitely. Therefore, utilizing Pachyderm on a cloud infrastructure impacts cost efficiency, in comparison to an on-premise high-performance computing (HPC) infrastructure. A notable feature that is supported by Lifebit, is cost-efficiency through low-priority instance utilization on a cloud provider, allowing for users to execute large-scale analyses at a reduced cost.

### Resource allocation

Pachyderm requires persistent RAM/CPU allocation for each pipeline within the Kubernetes cluster, even after a pipeline is successfully executed, which permits automatic pipeline triggering. Thus, an increased amount of compute resources (VM’s scaled up/out) may be required for specific pipelines, which also impacts cost efficiency.

### The ORCESTRA platform

Building on the strengths of the Pachyderm orchestration tool, we have developed ORCESTRA, a cloud-based platform for data sharing and processing of biomedical data based on automation, reproducibility, and transparency. ORCESTRA allows users to create a custom data object that stores molecular profiles, perturbation (chemical and radiation) profiles, and experimental metadata for the respective samples and patients, allowing for integrative analysis of the molecular and perturbation and clinical data (Figure 1). The platform utilizes datasets from the largest biomedical consortia, including 17 curated data objects containing genomics, pharmacological, toxicological, radiation and clinical data (Supplementary Table 1). The data objects can accommodate all types of molecular profile data, however, ORCESTRA currently integrate gene expression (RNA-sequencing, microarray), copy number variation, mutation, and fusion molecular data. For RNA-seq data, users can select a reference genome of interest, a combination of quantification tools and their respective versions, along with reference transcriptomes from two genome databases (Ensembl, Gencode) to generate custom RNA-seq expression profiles for all of the cell lines in the dataset. Therefore, each data object will be generated through a custom orchestrated Pachyderm pipeline path, where each piece of input data, pipeline, and output data option is tracked and given a unique identifier to ensure the entire process is completely transparent and reproducible. To ensure data object generation is fully transparent and that provenance is completely defined, each data object is automatically uploaded to Zenodo and given a public DOI, where the DOI is shared via a persistent webpage that possesses a detailed overview about the data that each DOI-associated data object contains and how it was generated. This includes publication sources, treatment sensitivity information and source, raw data source, exact pipelines parameters used for the processing tools of choice, and URLs to reference genomes and transcriptomes used by the tool(s). In addition, release notes are also provided where the number of samples, treatments, sensitivity experiments, and molecular profile data are tracked between versions of a dataset, allowing users to identify changes between each new data update that were released from the respective consortium and pushed to the platform. This metadata page gets automatically sent to each user via email, providing users with one custom page that hosts all of the information required to understand how the data object was generated. Therefore, all of the data used in the data object is shared in a transparent manner, where researchers can identify the true origins of all data used with confidence and effectively reproduce results.

**Figure 1.**
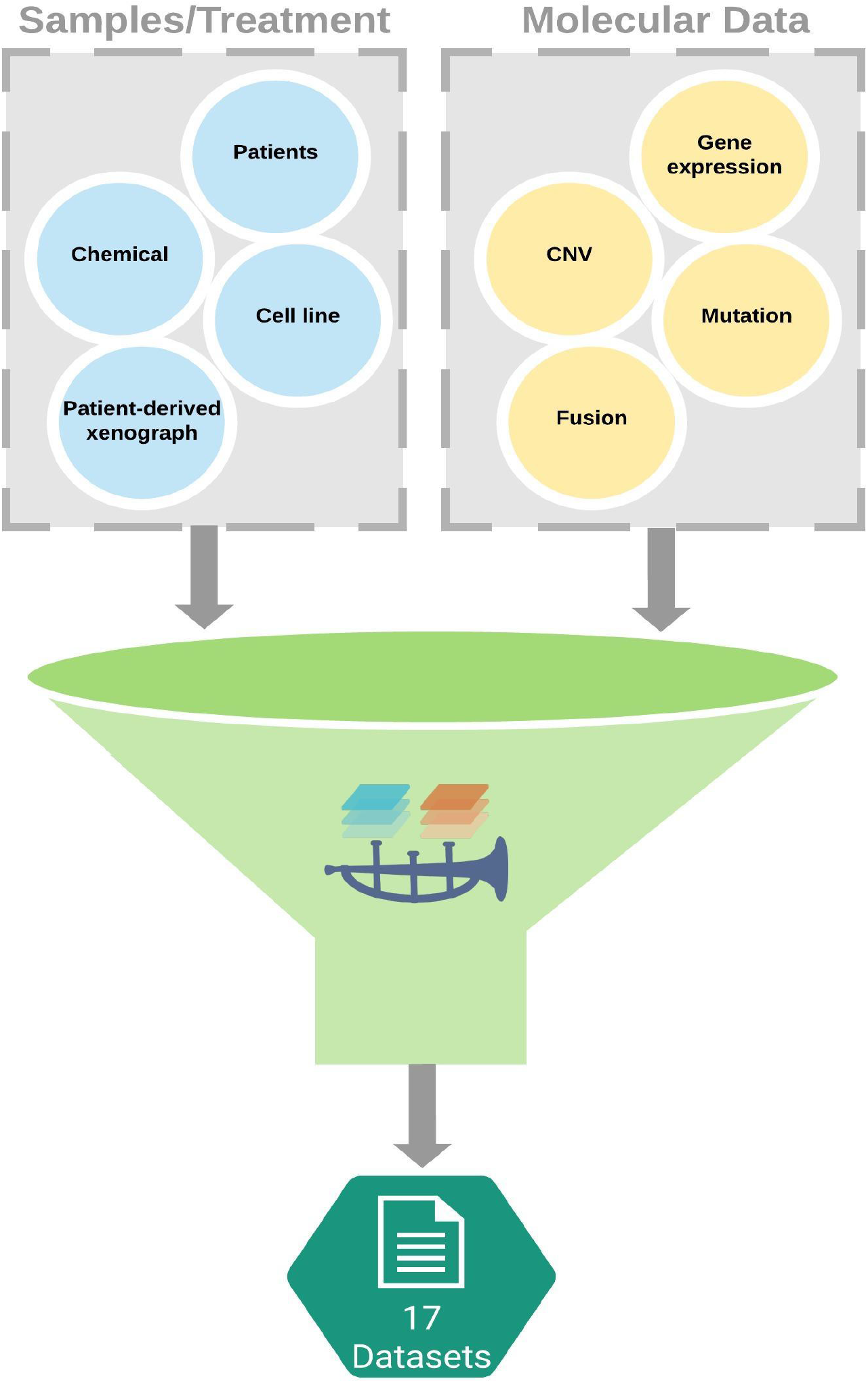
Summary of samples, treatments, and molecular profiles utilized for data object generation in ORCESTRA.

### Data object generation

*ORCESTRA* comes with a web application interface allowing users to interact with the data-processing and data-sharing layers. Users can search existing data objects in the “Search/Request’’ view by filtering existing data objects with the “data object Parameters’’ panel. Users can filter existing data objects by selecting datasets with associated drug sensitivity releases, genome references, RNA-seq transcriptomes, RNA-seq processing tools with respective versions, which associates with other respective DNA data types (mutation or CNV) and RNA data types (microarray or RNA-seq). Changes in the parameter selections trigger the web app to submit a query request to a MongoDB database which returns a filtered list of data objects (Figure 2). The data object table is then re-rendered with an updated list of data objects. This allows users to search through existing data objects to determine if a data object that satisfies users’ parameter selections already exists, preventing recomputation. Information about the datasets and tools used to generate a data object can be viewed by clicking on a data object name and navigating to its data object meta-data webpage. Users can obtain information such as associated publications, links to the raw drug-sensitivity and molecular profile data as well as a Zenodo DOI. In addition, the individual data object view provides users with the option to download the data object of choice directly from the view.

**Figure 2.**
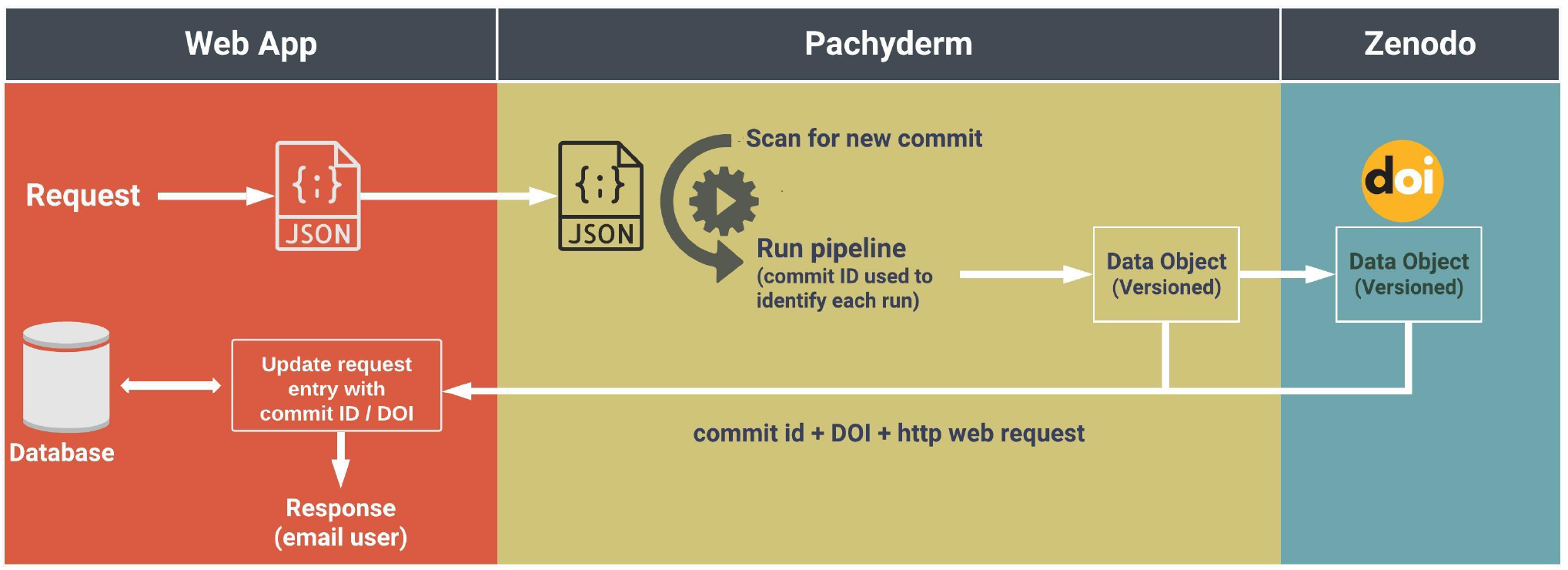
ORCESTRA web-application connectivity with data processing layer through commit ID scanning for user selected pipeline requests, and subsequent data object DOI tracking with MongoDB queries.

Users can request a customized data object in the “Search/Request’’ view by turning the “Request data object’’ toggle on. This action reconfigures the dropdown options in the “data object Parameters’’ panel to be in request mode, and displays, on the “Summary” panel, two text input fields for entering a custom name for the data object and a user’s email to receive a notification upon data object pipeline completion, with the accompanied Zenodo DOI and custom ORCESTRA meta-data page link. Pachyderm continuously scans for a new request from the web-app, which will automatically trigger the respective pipelines to build the custom data object, while storing a unique ORCESTRA ID, Pachyderm pipeline commit ID, and Zenodo DOI into the MongoDB database, which increases the level of data provenance and reproducibility, as each data object can be identified through three unique-identifiers after creation (Figure 2). The data object filtering process as described above continues to function as users select the request parameters, which displays existing data object(s) that satisfy users’ parameter selections. Upon selecting all the required parameters, the “Submit Request” button becomes active for users to submit the pipeline request.

### Data object metrics

The platform provides several usage metrics for users. These metrics can be accessed through “Home”, “Statistics’’ and “Request Status” views. The “Home” view provides an overview of currently available datasets, tools and references to generate data objects, most downloaded data objects, and a number of pending or in-process data object requests. The “Statistics’’ view provides a visualized data object popularity ranking, along with a plot of the number of cell lines, drugs, and genes for the canonical data objects, including intersection, which can be accessed by clicking the “View Statistics’’ button in the “Home” view. The “Request Status” view displays a tabulated list of data object requests that are either “pending” (the request has been submitted and saved, but has not been processed in Pachyderm), or “in-process” (the request has been submitted and is processed in Pachyderm).

### User accounts for data object tracking

The platform offers users the option to register for an account with a valid email address. Registered users are able to select existing data objects in the “Search/Request’’ view and save them as their “favorites’’ which can be accessed in the “User Profile” view. However, the web application keeps track of data object requests submitted by users based on their email addresses even without registration. These data objects are automatically added to a user’s favorite data objects and can be viewed in the “User Profile” view.

## DISCUSSION

The high-dimensionality, complexity, and scale of multimodal data present unprecedented challenges for researchers in the biomedical field, in regard to their ability to effectively manage, track, and process the data. The nature of heterogeneous and complex data negatively impacts data provenance, through incomplete or no accompaniment of metadata for a dataset, resulting in the uncertainty of a data lineage^24–26^. Because the granularity of metadata is a determinant of the value of a dataset^27^, it should provide a rich description of dataset content, following the FAIR data principles, which includes information about dataset origin, how it was generated, if there were any modifications that were made to it from precedent versions, and what these modifications were^21,28,29^. When the FAIR data principles are not met, issues with reproducibility in the biomedical sciences follow, where data are either not shared or results/estimates and claims cannot be checked for correctness. However, datasets published online, including ones that reside in repositories and from journals are often not accompanied with sufficient metadata^30^. In the field of genomics, issues with metadata often include mislabelling or misannotation of data (e.g., incorrect identification numbers), improper data characterization (e.g., mapping files to respective samples and protocols), and inconsistency in the way metadata are presented (nonuniform structure used across consortia)^31^. Provenance also extends to the computational workflows that are developed to process datasets^2^, as sharing relevant source code is often not provided^32^ along with relevant documentation about the workflow, such as in graphical user interface (GUI) based systems like Galaxy, affecting the ability to reproduce results^2^. In addition, data maintainers and consortia, such as the Cancer Cell Line Encyclopedia (CCLE)^33^ and the Genomics of Drug Sensitivity in Cancer (GDSC), often only process the dataset using one pipeline that they believe is the most suitable, without documenting supporting evidence as to why the chosen processing pipeline was selected over other competing ones in the field^34,35^. This issue is also present in other data types such as xenographic or metagenomics data, where the molecular data are processed and normalized using only one pipeline^20,36^. Therefore, only a single version of the dataset is released, which makes it difficult for other researchers to perform a diverse set of analyses that require the use of different processing pipelines on the dataset. A lack of provenance and utilization of a dataset in a single form affects transparency, expressing a need for sharing biomedical data in a reproducible manner. Lastly, it is important to note that datasets evolve and are therefore not static, as new data are added and respectively depreciated, which further highlights the need for transparent data sharing practices, especially at the file-level where updates can be easily identified.

There are multiple data portals created for accessing and sharing biomedical data, but with limitations in regard to reproducibility (Supplementary Table 2). Below, are sharing practices that are adopted across various data types, such as pharmacogenomics, toxicogenomics, radiogenomics, xenographic pharmacogenomics, and clinical genomics data:

### Pharmacogenomics

The Genomic Data Commons Data Portal (NIH/NCI GDC) hosts raw data for the Cancer Cell Lines Encyclopedia (CCLE) from the Broad Institute, including RNA, whole exome and whole genome sequencing data, allowing users to select and download the data type(s) of interest. Obtaining the data can be done through direct download or their GDC Data Transfer Tool by providing a manifest file that possesses the unique identifiers (UUID) of each file, which also allow users to locate the files again through the portal, along with their corresponding run, analysis, and experimental metadata. This is advantageous, as all the raw data (public and controlled access), for both datasets are located within one portal and can be accessed in an efficient manner. However, no release notes are provided for any data that are newly uploaded or modified to GDC, which makes it challenging for researchers to keep track of different versions of the dataset and ensure their analyses are reproducible. Moreover, the recent addition of new CCLE data (e.g., additional RNA-seq cell lines)^34^, is found on the European Nucleotide Archive (ENA), but not on GDC, resulting in data source inconsistency that becomes difficult to manage and follow for users. Current and previous versions of other CCLE data (i.e., annotation, drug response) are hosted on a Broad Institute portal, with no release notes or documentation present with each version, forcing researchers to manually identify changes within each file after every release. GRAY, a dataset generated by Dr. Joe Gray’s lab at the Oregon Health and Science University, has had three updates with raw data hosted on NCBI, with drug response and annotation data hosted on SYNAPSE, DRYAD, and/or the papers supplementary section^37–39^. In addition, drug response data can also be found on the LINCS data portal. Because each version of the dataset is associated with a different respective paper, the data are scattered among various repositories, which makes it challenging to keep track of each source, and for each source to ensure that the data remain readily available, as one failed link would make it difficult for a researcher to reproduce any results. However, for the GRAY dataset, NCBI provides detailed information about the methodology used for the experiments, SYNAPSE provides a wiki and contact source for the dataset and a provenance tracker for each file that is uploaded, and DRYAD stores each publications data as a package organized with subsequent descriptions to keep data organized. A prominent example in effective data sharing practices is DepMap (depmap.org)^40^, which provides a portal to download molecular and pharmacological data from a variety of consortia, with an interface that allows users to dive into the multiple data releases for a given dataset, which is accompanied by descriptive metadata such as associated publications and file-level descriptions. This provides users with the ability to download a dataset directly from a source, or combine them together to form a custom dataset, all while being able to compare different updates/versions in an interactive manner. However, the portal does not allow users to select different processing pipelines and lacks details regarding the pipelines used for some of the processed data hosted, such as molecular data (e.g., genomic tools used), which highlights a need for increased granularity in the portal.

### Toxicogenomics

The Life Science Database (LSDB) Archive is a database that hosts datasets by Japanese researchers (https://dbarchive.biosciencedbc.jp/), such as the TG-GATE toxicogenomics dataset^41^. The database provides rich metadata for users such as a DOI and clickable sections that provide granular details about each file in the dataset, which includes a description of the file contents and file attributes (e.g., data columns and respective descriptions for each column). In addition, the database allowed for TG-GATE to provide a timeline of updates to the dataset, where data corrections are posted with accompanying corrected files and a description of the update, which allows maintainers to be transparent with users about the dataset lineage. However, even though the maintainers for TG-GATE have indicated that the dataset was updated, detailed file-level changes are not provided, along with the processing pipelines and/or information regarding how the data was generated/processed into their resulting formats.

### Xenographic and Radiogenomics

The largest datasets for patient-derived tumor xenograft and radiogenomics studies are available through supplementary materials attached to their scientific publications^19,36^. These supplementary data provide users with information about the methods used to generate the data, however access is dependent on the journal itself, which raises issues regarding the potential of broken data links. In addition, the amount of data that can added to a publication via a supplementary section may be limited due to journal restrictions, which increases the likelihood of files being distributed across other data sharing platforms (e.g., SYNAPSE), increasing the difficulty in locating and keeping track of dataset updates, or resulting in a reliance of contacting authors to obtain a complete dataset that cannot be otherwise shared via the journals web interface.

### Clinical Genomics

Over the years, clinical genomics data has been stored and shared across a wide range of consortia such as NCBI (GEO/EGA) and/or as supplementary material to a publication. However, this inconsistency has led to the development of several data compendias to consolidate the data for transparent mining/managing, sharing, and analysis, such as Oncomine^42^, MultiAssayExperiments R package for multiple experimental assays^43^, and curatedData R packages for molecular profile analysis^44^. In addition, the MetaGx R packages were developed to allow users to retrieve a compendium of transcriptomic data and standardized metadata from a wide array of studies and cancer types (pancreas, breast, ovarian), allowing for integrative analysis of the data for biomarker discovery^20^.

In conclusion, the ORCESTRA platform provides a new paradigm for sharing ready-to-analyze multimodal data while ensuring full transparency and reproducibility of the curation, processing and annotation processes. ORCESTRA provides the data provenance and versioning tools necessary to maximize reusability of data, a cornerstone of Open Science.

## METHODS

In order for the platform to be as transparent as possible, it harnesses an architecture with three distinct layers that not only works independently to process and interpret precedent data, but also have the capacity to scale (Figure 3).

**Figure 3.**
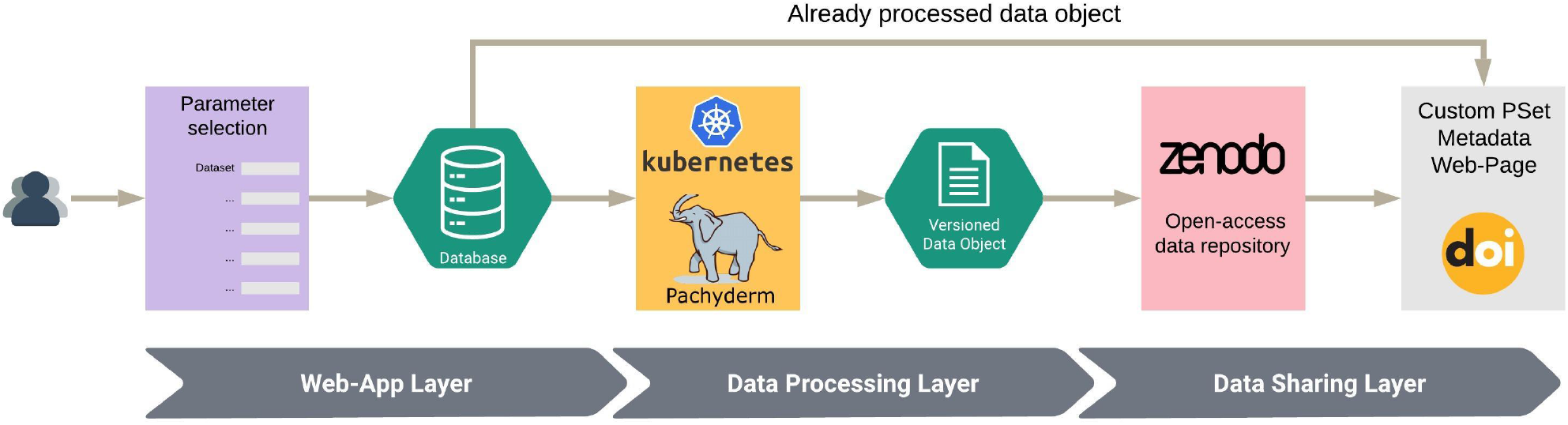
ORCESTRA framework layers for pipeline selection, data object generation, and DOI sharing with a custom metadata webpage.

### Web-app layer

The first layer contains the web application which was developed using a Node.js API and React front-end with MongoDB as a database. The layer provides the user with an interaction point to the *ORCESTRA* platform, allowing users to first select the data type they wish to explore, either Pharmacogenomics; Toxicogenomics; Xenographic pharmacogenomics; Radiogenomics; or Clinical genomics. They can then search for existing data objects, request a new data object by entering pipeline parameters, view data object request status, and register a personal account to save existing data objects of choice.

### Data-processing layer

The second data-processing layer encompasses a Kubernetes cluster on Microsoft Azure that hosts Pachyderm, which utilizes Docker images for running R-packages. All of the RNA-seq raw data have been pre-processed with Kallisto and Salmon Snakemake pipelines using an HPC environment, and subsequently pushed to assigned data repositories on Pachyderm, allowing for specified selection from the web-app (transcriptome and tool version). Microarray, cnv, mutation, and fusion data are either processed directly with Pachyderm due to low compute requirements or aggregated into the data objects from public sources. The Pachyderm pipelines aggregate repositories that host data generated on an HPC environment or on GitHub into a Docker image that builds a data object based on user specifications (e.g. RNA-seq data processed by Kallisto v.0.46.1, inclusion of only CNV data) (Figure 4). The GitHub hosted files can be viewed at the file-level for changes and edited which automatically triggers the Pachyderm pipeline with the new modifications to produce a new data object. A unique feature of Pachyderm is the prevention of re-processing computed data, such as where an update of RNA-seq annotations will not trigger the re-processing of thousands of drug response data, which reduces computation time.

**Figure 4.**
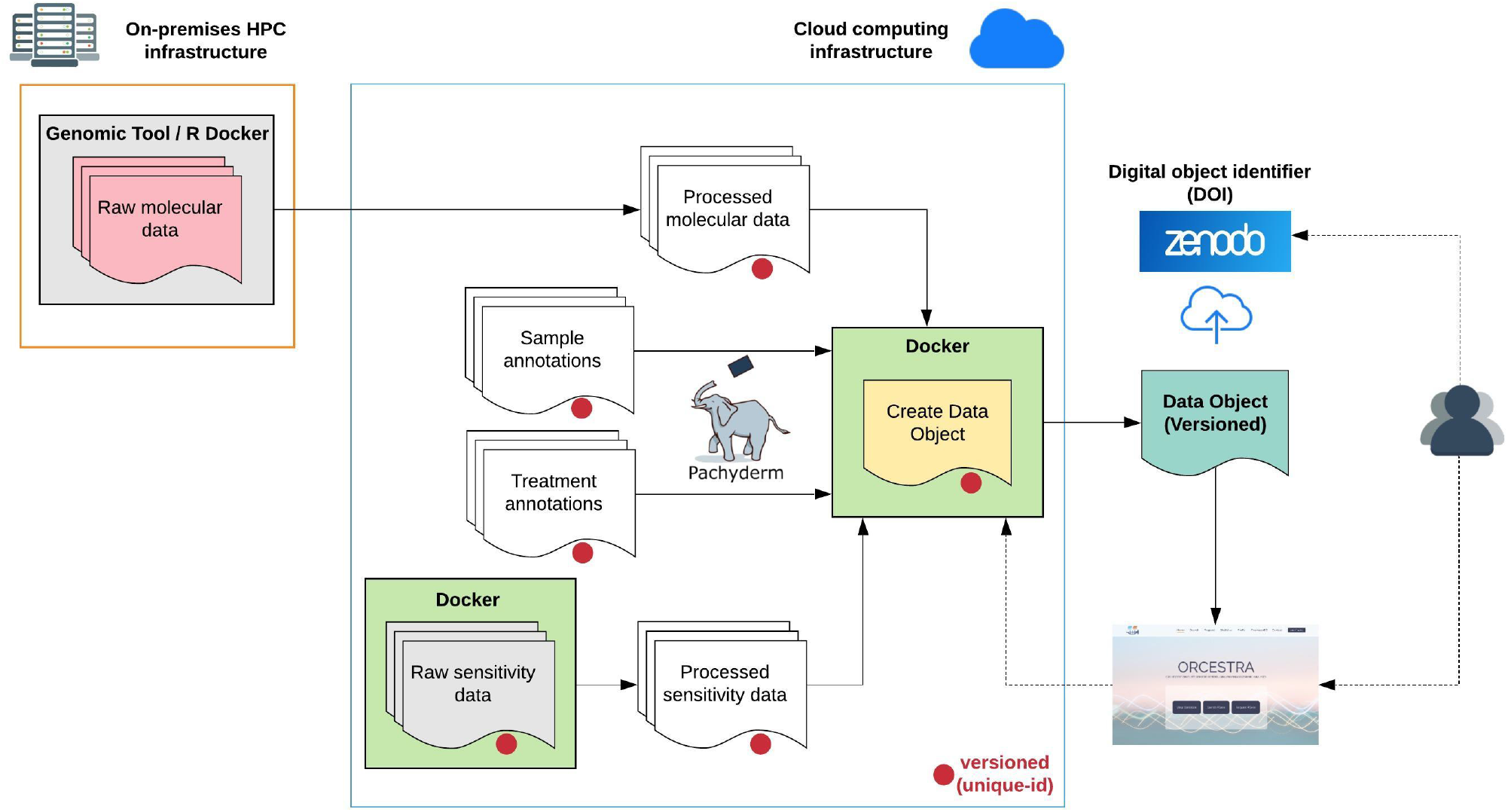
Cloud-based deployment of ORCESTRA data processing layer with automatically versioned data using Pachyderm and sharing of generated data objects through Zenodo via a persistent identifier (DOI).

### Data sharing layer

Each generated data object enters the third data-sharing layer where the data object gets automatically uploaded to an online data-sharing repository known as Zenodo, with a DOI so that the data object can be given a persistent location on the internet to be uniquely identified. The generated DOI is then associated with a custom meta-data web page that is generated based on the contents of the data object.

### Code availability

All of the code is publicly available on GitHub via the Apache 2.0 license: https://github.com/BHKLAB-Pachyderm

## Data availability

All of the data are publicly available on ORCESTRA (orcestra.ca) via dedicated documented webpages, which include respective Zenodo links for each data object generated.

## Acknowledgements

This project is supported by the Canadian Institutes of Health Research (CIHR), under the frame of ERA PerMed.

