## Supplementary Table 1 for "Orchestrating and sharing large multimodal data for transparent and reproducible research"

| Category | Dataset name | No. of samples | No. of perturbations | Release date/version |
|---|---|---|---|---|
| Pharmacogenomics (in vitro) | GRAY | 73 | 89 | 2013 |
| | GRAY | 74 | 107 | 2017 |
| | FIMM | 50 | 52 | 2016 |
| | GDSC1 | 1104 | 343 | 2020(v1-8.2) |
| | GDSC2 | 1104 | 190 | 2020(v2-8.2) |
| | gCSI | 747 | 16 | 2017 |
| | UHNBreast | 84 | 8 | 2019 |
| | CTRPv2 | 887 | 544 | 2015 |
| | CCLE | 1094 | 24 | 2015 |
| | GDSC1 | 1104 | 303 | 2019(v1-8.0) |
| | GDSC2 | 1104 | 169 | 2019(v2-8.0) |
| Pharmacogenomics (in vivo) | PDXE | 277 | 62 | 2015 |
| Radiogenomics | Cleveland | 540 | 1 | 2016 |
| Toxicogenomics | DrugMatrix | 1 | 126 | 2019 |
| | EMEXP2458 | 2 | 6 | 2010 |
| | Open TG-GATEs | 2 | 152 | 2015 |
| Clinical Genomics | MetaGxPancreas | 1792 | NA | 2019 |

**Supplementary Table 1.** Summary of data types with respective dataset names, samples, perturbations, and releases available for exploration through ORCESTRA.
