## Supplementary Table 2 for "Orchestrating and sharing large multimodal data for transparent and reproducible research"

| Data Portal | Notable Functionalities | Expected Functionalities |
|---|---|---|
| SYNAPSE | Wiki integration<br>Data provenance tracker<br>Dataset discussion board | Track updates to a dataset at the file level |
| Broad Institute (CCLE) | Easy access of current, previous, and legacy data versions<br>Display of data by data-type<br>(e.g. pharmacological profiling, mRNA expression) | Display of data processing tools, and their versions, with all accompanying files (e.g. transcriptome) |
| DRYAD | Simplistic navigation<br>Dataset metrics<br>Respective publication download | Direct access to all processing pipelines utilized |
| NCBI | SRA sequencing information with additional sample metadata | Option of downloading processed data through multiple methods/pipelines |
| CancerRxGene (GDSC) | Direct integration of cell line and drug metadata<br>Direct links to raw data hosted on other portals<br>Description of changes to each drug sensitivity data release | |

**Supplementary Table 2.** Common data portals for sharing genomics data with notable and expected functionalities for transparent and reproducible processing, analysis, and interpretation.
